# MitoNGS: an online platform to analyze fish metabarcoding data in high-resolution

**DOI:** 10.1101/2025.09.29.679108

**Authors:** Tao Zhu, Yukuto Sato, Tsukasa Fukunaga, Masaki Miya, Wataru Iwasaki, Susumu Yoshizawa

## Abstract

Environmental DNA (eDNA) metabarcoding has become a powerful tool for assessing fish biodiversity in aquatic ecosystems. However, accurate species-level identification remains challenging due to incomplete and contaminated reference databases, as well as ambiguous taxa sharing identical barcode sequences. Here, we present MitoNGS, a next-generation platform that succeeds the widely used MiFish pipeline, designed for high-resolution analysis of fish metabarcoding data. MitoNGS addresses these challenges by incorporating more comprehensive references including non-fish species and detailed annotations of heterospecific regions. Additionally, it introduces the “species group” strategy in conjunction with environmental habitat and geographic occurrence data to resolve ambiguous taxa. Furthermore, MitoNGS expands the functionalities of the legacy MiFish pipeline. It can analyze data from any mitochondrial markers and from Nanopore sequencing platforms. MitoNGS demonstrated excellent performance on our testing datasets from diverse locations, markers and sequencing platforms. MitoNGS offers a user-friendly, web-based solution for fish detection, biodiversity monitoring, conservation research, and bioresource management. MitoNGS is freely available via https://mitofish.aori.u-tokyo.ac.jp/mito-ngs.

## 1 Introduction

Metabarcoding of environmental DNA (eDNA) targeting mitochondrial genes has emerged as a powerful, efficient, and non-invasive approach for monitoring fish diversity in aquatic environments. The MitoFish database and MiFish pipeline (Iwasaki et al., 2013; Sato et al., 2018; Zhu, Sato, Sado, Miya, & Iwasaki, 2023) were developed to analyze fish eDNA metabarcoding data and have been widely used and cited in recent years (Jiang et al., 2025; Tsuji, Takahara, Doi, Shibata, & Yamanaka, 2019; Xiong et al., 2022; Yang et al., 2024; Zhu et al., 2023). However, several challenges remain to enhance the benefits of fish eDNA research, including the application of alternative markers, improved resolution, a multi-sample strategy, and portable sequencing.

The online MiFish pipeline was specifically designed to analyze sequencing data from the MiFish primers (Miya et al., 2015), including MiFish-U and MiFish-E within the 12S rRNA gene for universal fish species and elasmobranchs, respectively. Although these two primers have been proven to be the most efficient and widely used (Xiong et al., 2022), other markers such as Teleo (Valentini et al., 2016) or Riaz (Riaz et al., 2011), are still being utilized in various researches. Although a standalone version of the MiFish pipeline based on a command line interface was established two years ago (Zhu et al., 2023), it requires creating and formatting customized reference databases, which is less convenient than the online platform. An updated platform with the capability to handle any mitochondrial markers is essential for expanding fish eDNA research.

The resolution of taxonomy identification remains a critical challenge in metabarcoding. Species serves as the fundamental unit of biological communities, and ideal metabarcoding outcomes should solely present a comprehensive list of existing species in the samples, which is often impractical. Two primary obstacles hinder species-level identification: incomplete reference databases and ambiguous barcode sequences. Incomplete reference databases, such as those derived from endemic species, lack sequences of invasive and off-target species, resulting in incorrect matching, particularly in instances of low matching identities. The NCBI-nt database (also known as GenBank) (Sayers et al., 2025) includes sequences from all organisms and was directly utilized as a reference database in several fish eDNA research projects (Gao et al., 2025; Plewnia, Krehenwinkel, & Heine, 2025; Rossouw, von der Heyden, & Peer, 2025). However, it has been argued to contain errors that could substantially impact the results (Steinegger & Salzberg, 2020). On the other hand, ambiguous barcode sequences arise when a single amplicon sequence variant (ASV) matches multiple species with high (99% or even 100%) and identical matching identities. Most algorithms assign lowest common ancestors (LCAs) to such ASVs (Griffiths et al., 2025; Polanco F et al., 2025), which might be overly ambiguous to interpret, as LCAs typically contains numerous other species not belonging to the top-hits. The current version of the MiFish pipeline (Sato et al., 2018; Zhu et al., 2023) randomly selects one species, marks it as low confidence, and additionally presents an alternative one. Nevertheless, it remains uncertain whether other alternative species exist. Several studies have assigned a virtual “group” containing solely the top-hit species to the ASV (Poiesz et al., 2025; Rehill et al., 2024). Such groups have also been referred to “species complex” or “species consensus”. We consider this approach appropriate for reflecting the biological origin of ambiguous ASVs and designate them as “species groups” in subsequent sections. Although species within the same group have identical barcoding sequences, their environmental habitats such as water depth, salinity, temperature, and geographic distribution, may not be identical. Several studies have applied geographic occurrence information to facilitate the filtering of species groups (Anil et al., 2025; Kurniawan et al., 2025; Rehill et al., 2024; Tibone et al., 2025). The combination of a complete and curated reference database (MitoFish), an off-target database, species groups, environmental habitat data and geographic occurrence data will be instrumental in enhancing the taxonomy resolution of metabarcoding.

Incorporating positive controls, negative controls, and biological or technical replicates serves as an effective strategy to mitigate or eliminate the impact of cross-contamination during eDNA metabarcoding. Several analysis pipelines such as VTAM (González et al., 2023) and AMPtk (Palmer, Jusino, Banik, & Lindner, 2018), have been developed to address these specialized samples. While the current MiFish pipeline can analyze multiple samples, they are treated uniformly without considering their distinct roles. By incorporating such specialized samples, it will contribute to the reduction of false positive results.

Furthermore, portable Nanopore sequencing devices such as MinION have been successfully applied to on-site eDNA sequencing (Krehenwinkel, Pomerantz, & Prost, 2019). This would be a significant advancement in enabling real-time monitoring of fish diversity. Notably, Nanopore sequencing has the capability to sequence long amplicons, thereby enhancing the distinction between closely related species. However, data analysis of the noisy Nanopore reads is challenging and typically incurs substantial computational resources. Currently, there are limited eDNA analysis pipelines capable of handling Nanopore reads. The development of an online platform dedicated to analyzing Nanopore raw reads would undoubtedly streamline the entire metabarcoding process. Users would only require a laptop or tablet connected to the internet, enabling the upload of sequencing results produced by MinION to the online analysis platform for rapid species composition identification.

Here, we have developed MitoNGS, the successor of the MiFish pipeline, to analyze fish eDNA metabarcoding data from any mitochondrial markers and multiple sequencing platforms such as Illumina and Nanopore. MitoNGS utilizes MitoFish and a newly established background database as references to ensure completeness and accuracy. By employing multiple strategies, it generates high-resolution taxonomy annotation results at the species-level or species groups. MitoNGS is publicly available via https://mitofish.aori.u-tokyo.ac.jp/mito-ngs, while the URL of legacy MiFish pipeline https://mitofish.aori.u-tokyo.ac.jp/mifish will be automatically redirected to MitoNGS.

## 2 Methods

### 2.1 Reference database

The reference database MitoFish comprises mitochondrial sequences, validated taxonomy labels, environmental habitat and geographic occurrence data. These data are updated monthly, and historical snapshots are publicly available via https://mitofish.aori.u-tokyo.ac.jp/download/.

#### 2.1.1 Sequences and gene annotations

Fish mitochondrial sequences were retrieved from the GenBank database (https://www.ncbi.nlm.nih.gov/nuccore) using the following searching strategy: txid7742[Organism:exp] NOT txid32523[Organism:exp] AND mitochondrion[filter] AND ddbj_embl_genbank[filter] NOT uncultured[filter] NOT UNVERIFIED[title] NOT UNVERIFIED_ASMBLY[title] NOT UNVERIFIED_ORG[title] NOT CON[Division]. Downloaded sequences were merged into a single file in FASTA format, combining identical sequences to reduce redundancy while preserving their accession IDs in the sequence titles. The majority of sequences originate from single mitochondrial genes, and their names were standardized according to Table 1. Sequences containing multiple mitochondrial genes (i.e., complete or partial mitogenomes), and those lacking any gene annotations, were re-annotated using MitoAnnotator (Iwasaki et al., 2013; Zhu & Iwasaki, 2023).

**Table 1.** Uniformed names of mitochondrial genes to make further analysis convenient.

| Uniformed names | Description | Alternative names* |
| --- | --- | --- |
| rRNA genes |  |  |
| 12S rRNA | 12S ribosomal RNA | rrnS, rrn12, rns, rrnaS, s-rRNA, SSU, ... |
| 16S rRNA | 16S ribosomal RNA | rrnL, rrn16, rnl, rrnaL, l-rRNA, LSU, ... |
| protein coding genes |  |  |
| ND1 | NADH dehydrogenase subunit 1 | NAD1, NADH1, NADH-I, ... |
| ND2 | NADH dehydrogenase subunit 2 | NAD2, NADH2, NADH-2, ... |
| COXI | cytochrome c oxidase subunit 1 | COX1, COI, CO1, COX-1, ... |
| COXII | cytochrome c oxidase subunit 2 | COX2, COII, CO2, COX-II, ... |
| ATPase8 | ATP synthase F0 subunit 8 | ATP8, apt8, MTATP8, stp8, ATPase-8, AT8, ... |
| ATPase6 | ATP synthase F0 subunit 6 | ATP6, MTATP6, ATPase-6, AT6, ATPase-6, ... |
| COXIII | cytochrome c oxidase subunit 3 | COX3, COIII, CO3, COX-III, ... |
| ND3 | NADH dehydrogenase subunit 3 | NAD3, NADH3, NADH-3, ... |
| ND4L | NADH dehydrogenase subunit 4L | NAD4L, NADH4L, NADH-4L, ND41, NA4L, ... |
| ND4 | NADH dehydrogenase subunit 4 | NAD4, NADH4, NADH-4, ... |
| ND5 | NADH dehydrogenase subunit 5 | NAD5, NADH5, NADH-5, NND5, MD5, ... |
| ND6 | NADH dehydrogenase subunit 6 | NAD6, NADH6, NADH-6, ... |
| Cytb | cytochrome b | MT-CYB, cyto B, Cyt-B, CYBT, ctb, cob, Cb, ... |

\*Names are case-insensitive. For example, there are cases that COXI is written as COI or coi.

#### 2.1.2 Taxonomy validation of sequence regions

It is crucial to validate the taxonomy labels in reference sequences to prevent false positive metabarcoding outcomes. Recent studies have confirmed the presence of chimeric sequences in GenBank, wherein certain regions within a single sequence originate from heterospecific contaminations, while the remaining regions are homospecific and correctly labeled (Claver, Canals, de Amézaga, Mendibil, & Rodriguez-Ezpeleta, 2023; Sangster & Luksenburg, 2020). In this perspective, mislabeled reference sequences (X. Li et al., 2018) can be regarded as those containing 100% (or near 100%) heterospecific regions. The primary factor is whether the heterospecific regions overlap with the metabarcoding region, irrespective of the remaining regions. Consequently, we conducted the taxonomy validation on subsequence regions rather than full-length sequences. According to Sangster and Luksenburg (2020)’s research, heterospecific regions in fish’s mitochondrial genes can be as short as approximately 100bp. Therefore, we split all the 13 mitochondrial protein-coding genes and two mitochondrial rRNA genes into chunks of 100bp length. If the final chunk is shorter than 100bp, it was designated as the final 100bp of the gene sequence. For each chunk region, taxonomy validation was conducted based on the taxonomy and author information of other homologs with a similarity greater than 99%. Regions with homologs of the same species from different authors were considered homospecific, while regions with only homologs of different species from multiple authors were suggested as heterospecific (Figure 1).

**Figure 1.**
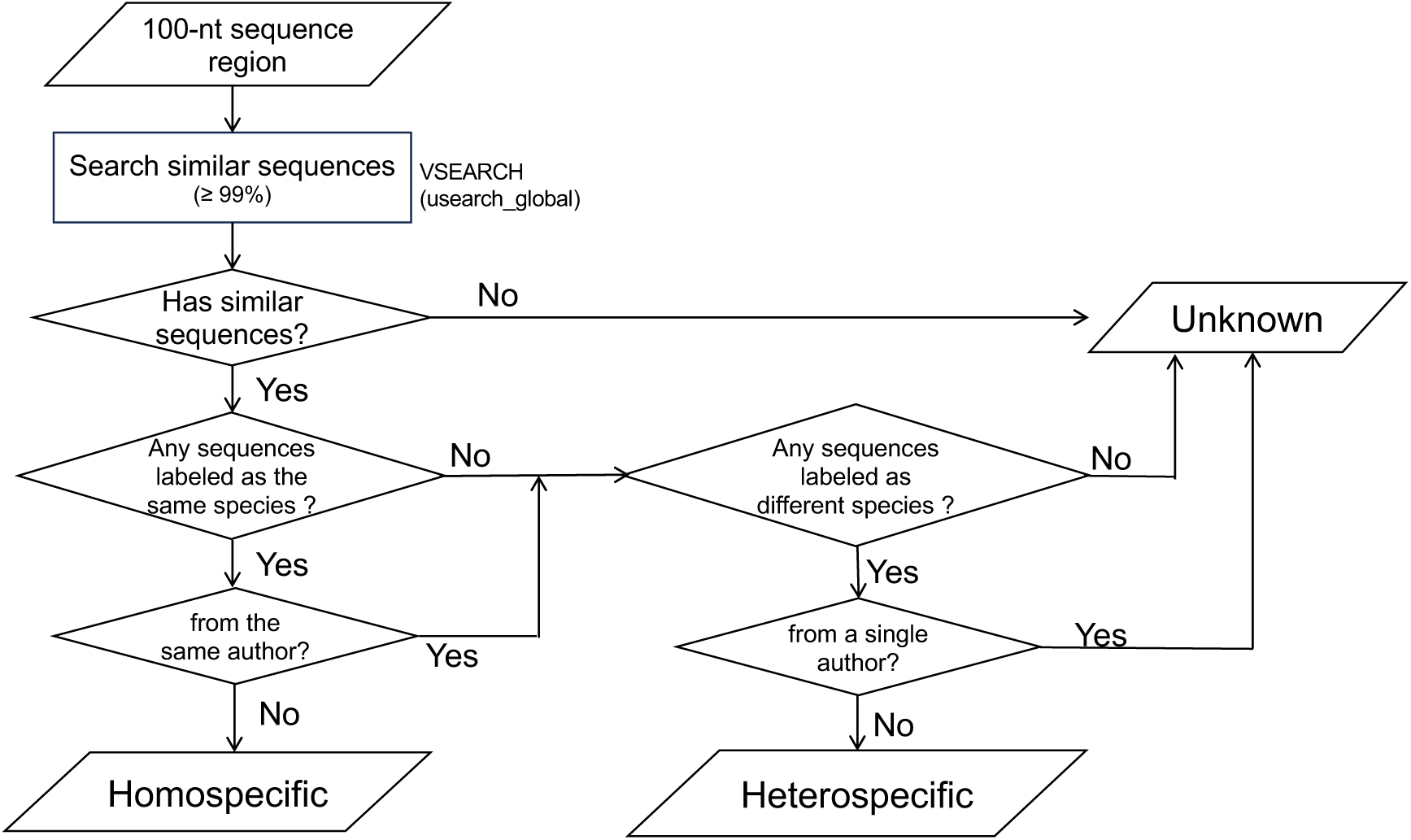
Workflows of judging homospecific and heterospecific mitochondrial gene region. Heterospecific and unknown regions will undergo validation monthly, with updates to the reference database.

#### 2.1.3 Environmental habitat data

Environmental habitat data for all fish species were sourced from rFishBase (Boettiger, Lang, & Wainwright, 2012). Three primary habitat features, “DemersPelag”, “Salinity”, and “Climate Zone” (Table 2), were selected as they encompass the fundamental aspects of aquatic environments. Notably, these features are available for over 99% of species in rFishBase. Since the species names in the reference database are from NCBI Taxonomy database, we performed a name mapping to FishBase via https://www.fishbase.org/tools/upload/CheckName.php.

**Table 2.** Environmental habitat data used in the reference database and MitoNGS.

| Type | Values |
| --- | --- |
| DemersPelag | bathydemersal<br>bathypelagic<br>benthopelagic<br>demersal<br>pelagic<br>pelagic-neritic<br>pelagic-oceanic<br>reef-associated |
| Salinity* | freshwater<br>brackish water<br>saltwater |
| Climate zone* | subtropical<br>temperate<br>tropical<br>boreal<br>deep-water<br>high altitude<br>polar |

\*One species might have multiple values.

#### 2.1.4 Occurrence data

Occurrence data with geographic coordinates for all fish species were obtained from Global Biodiversity Information Facility (GBIF, https://www.gbif.org). To minimize redundancy, occurrence data within the same species is deduplicated based on geographic coordinates (latitude and longitude) within a tolerance of 1 degree. Similarly, we conducted a name mapping from the NCBI Taxonomy database to GBIF via https://www.gbif.org/tools/species-lookup.

### 2.2 Background database

A background database was established to filter metabarcoding sequences originating from off-target amplification of non-fish species or cross-contamination from other metabarcoding experiments. It consists of two parts, non-mitochondrial genes, and mitochondrial genes of non-fish species. The former includes sequences of the 16S rRNA gene of prokaryotes and the 18S rRNA gene of eukaryotes obtained via SILVA v138.2 (Quast et al., 2012), and the internal transcribed spacer (ITS) of eukaryotes obtained via UNITE v10.0 (Abarenkov et al., 2024). The latter part includes all the 13 mitochondrial protein coding genes and two mitochondrial rRNA genes (12S rRNA and 16S rRNA) obtained via MIDORI2 v264 (Leray, Knowlton, & Machida, 2022), with sequences from fish species excluded by our local scripts. Since the background database serves as a filtering tool rather than a precise identification method for non-fish species, in order to reduce redundancy and speed-up the search, sequences were clustered using VSEARCH v2.30.0 (Rognes, Flouri, Nichols, Quince, & Mahé, 2016) at 97% identity, and the centroid for each cluster was kept in the background database.

### 2.3 Workflow of taxonomy identification

The taxonomy identification workflow comprises two distinct phases: the generation of amplicon sequence variations (ASVs) from raw sequencing reads (Figure 2), and the subsequent taxonomy annotation of ASVs (Figure 3).

**Figure 2.**
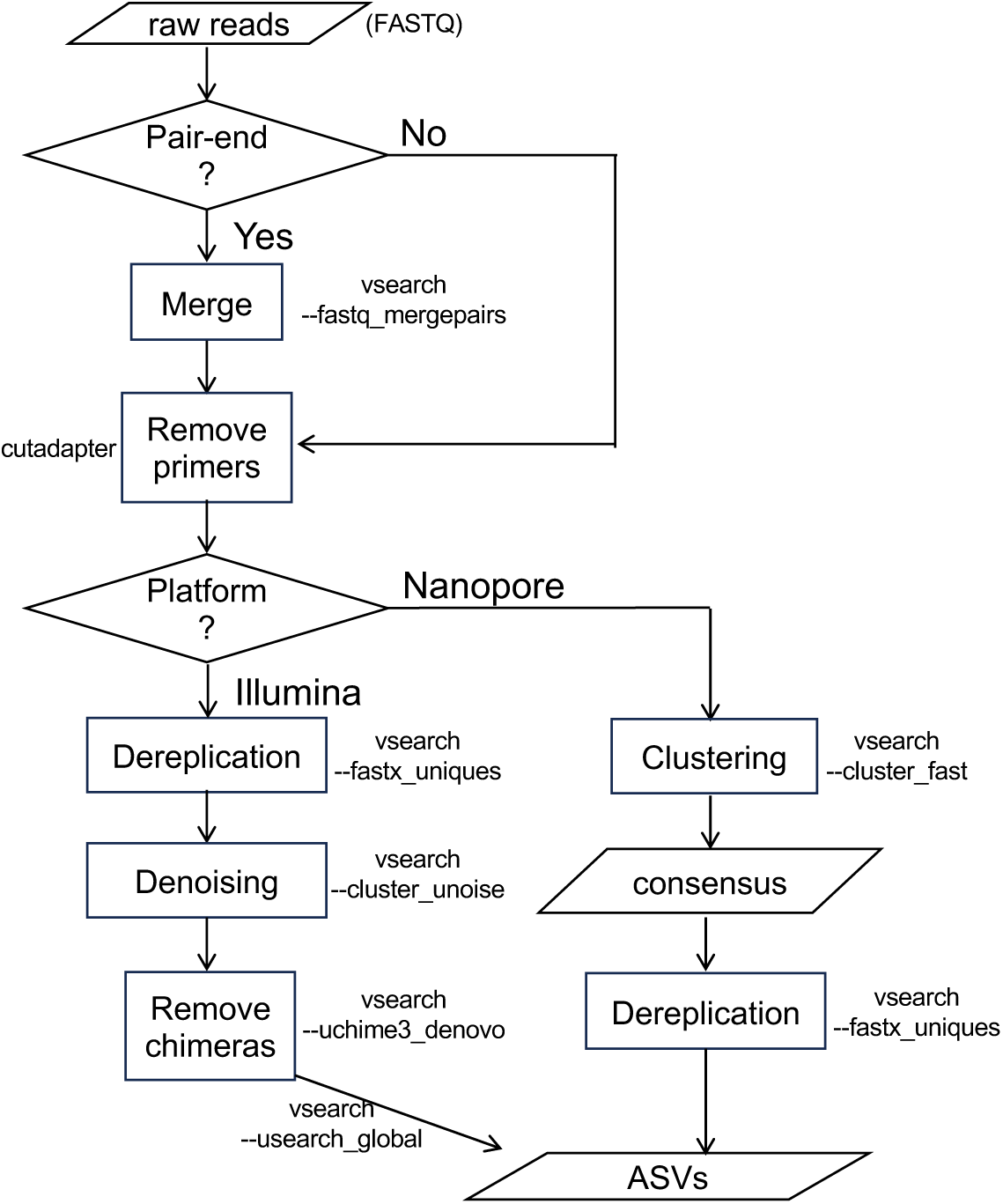
Workflows of generating amplicon sequence variants (ASVs) from raw sequencing reads.

**Figure 3.**
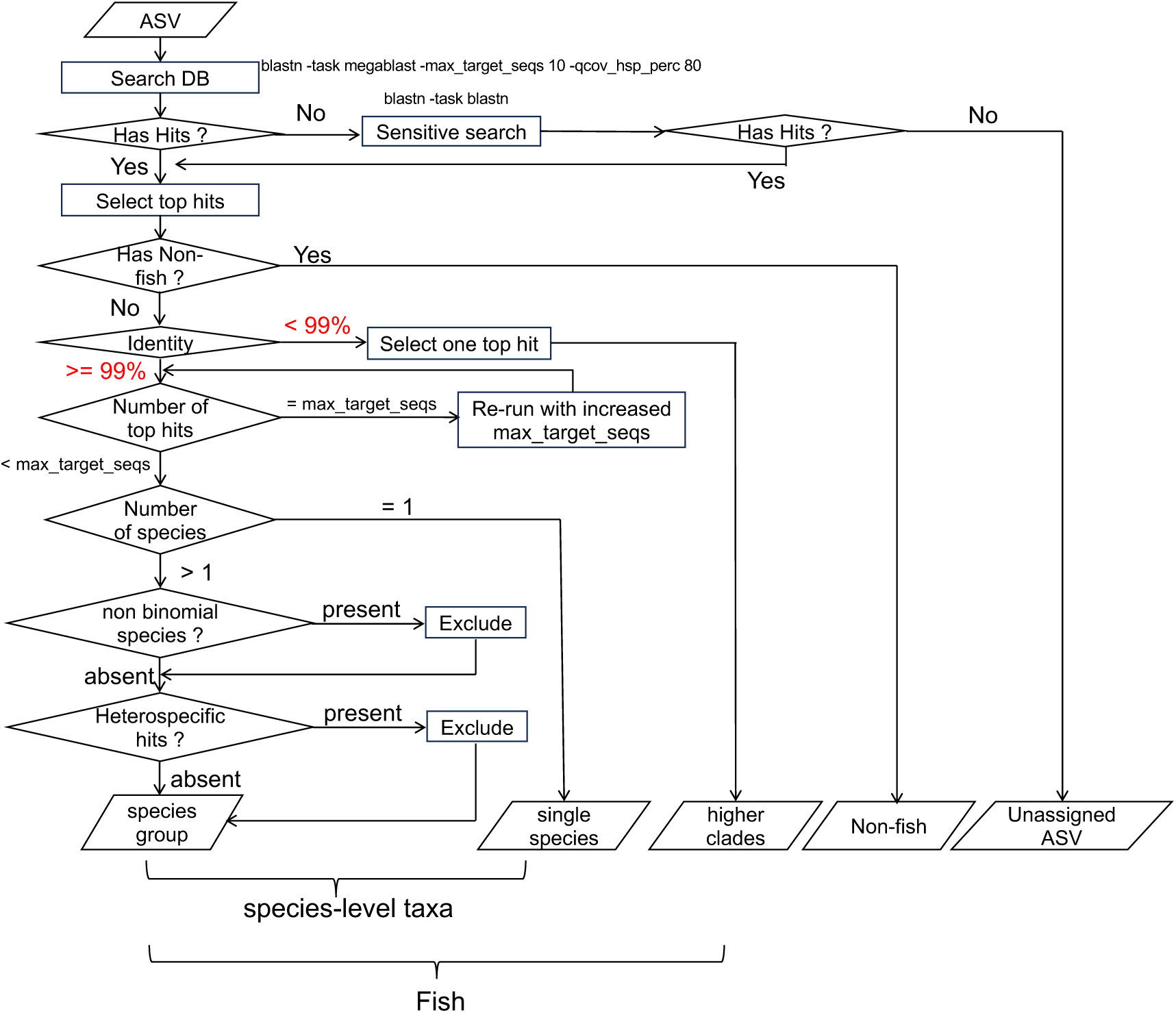
Workflows of taxonomy annotation of amplicon sequence variants (ASVs). 99% is the similarity threshold of annotating ASVs to species level, and this value is adjustable by users.

#### 2.3.1 ASV generation

The process of generating ASVs from reads of second-generation sequencing platforms, such as Illumina, is similar to the legacy MiFish pipeline (Zhu et al., 2023) except that we replaced USEARCH (Edgar, 2016) with the open-source VSEARCH v2.30.0 (Rognes et al., 2016) for denoising, as the latter is more efficient in handling large datasets. For noisy reads generated by third-generation sequencing platforms, such as Nanopore, we utilize the cluster_fast command in VSEARCH to cluster the noisy reads at a 97% identity threshold, employing penalties for gap openings set to “4I/2E” (recommended by VSEARCH developers at https://groups.google.com/g/vsearch-forum/c/qs_9Zo6cc8c/m/jlyFZHNjDwAJ), similar to the default behavior of Minimap2 (H. Li & Birol, 2018). The consensus sequence of each cluster is then used as an ASV for taxonomy annotation.

#### 2.3.2 Taxonomy annotation

In MitoNGS, the default similarity threshold for assigning ASVs to species level is set at 99%, which can be adjusted by users. BLASTN v2.16.0 is employed to search for similar sequences of ASVs in the reference and background database with parameters “-task megablast -max_target_seqs 10 -max_hsps 1 -qcov_hsp_perc 80”. For each ASV, the top hits with the highest score are retained for further analysis. If the number of top hits reaches 10 with identical highest score and their identities are ≥ 99%, indicating an unsaturated search, a recursive search method is employed. This method increases the value of “-max_target_seqs” by 50 during each iteration until a hit with a lower score is encountered. For ASVs that do not yield any hits, the parameter “-task megablast” is modified to “-task blastn”, which offers greater sensitivity but incurs a slower processing time, and the search is repeated. These strategies ensure that all ASVs receive their top hits from either reference or background database in a timely and exhaustive manner.

In the subsequent phase, each ASV was assigned to the species designated in the top hits. For ASVs originating from fish species and exhibiting an identity ≥ 99%, if multiple species coexist in the best hits, they undergo a filtering process through two additional steps. Firstly, non-binomial species, when coexisting with binomial species, are excluded because the former are not yet formally described and named. Secondly, species from which hits located in heterospecific regions were excluded due to insufficient evidence for the labelled taxonomy. If multiple species pass through the filtration process, they are kept as a “species group” that remains indistinguishable under this metabarcoding marker. Users retain the ability to perform further filtrations based on the environmental habitats or geographic occurrences, depending on the research content.

Finally, the taxonomy composition of the entire sample is categorized and presented in three distinct tables: species-level taxa, higher-level taxa and non-fish taxa. For species-level taxa, abundances are summarized by aggregating the read counts of all ASVs within the same species. While for higher-level and non-fish taxa, all ASVs are presented separately, accompanied by the MD5 checksum of the sequence.

#### 2.3.3 Controls and replicates

MitoNGS has adapted multi-sample analysis by presenting results in a table format, where rows represent species (including undistinguishable groups) and columns represent samples, including biological/technical replications, as well as positive/negative controls (Figure 4). Species detected solely in a single replicate or detected in controls are marked with specific symbols (Figure 4) to prompt users for manual inspection and filtration. Tables can be exported in CSV format for further analysis.

**Figure 4.**
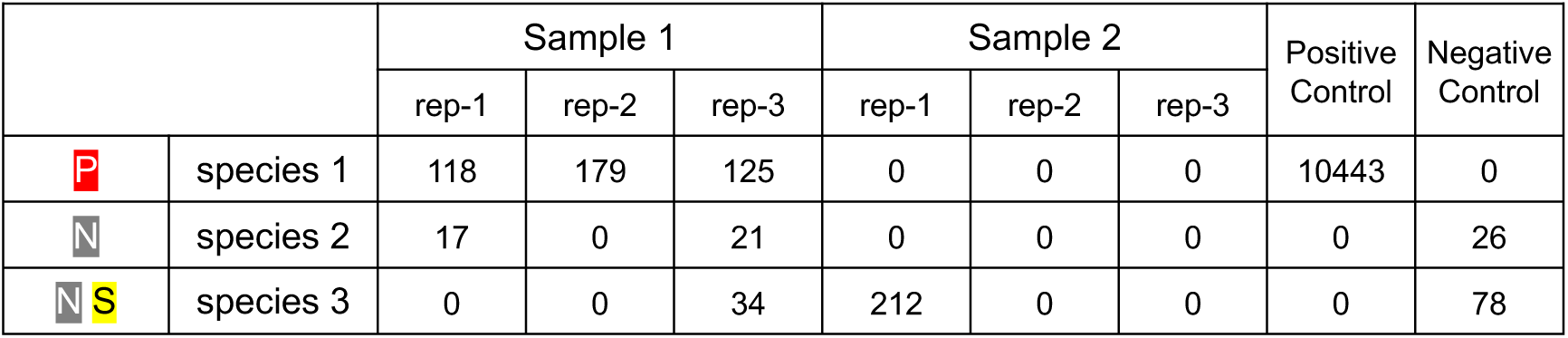
The structure of taxonomy identification results displayed in MitoNGS for multiple samples. Samples (including replications and controls) are arranged in columns, while species are positioned in rows. The abundance of each species is indicated by the count of reads in the corresponding table cells. The badge “P” signifies the detection of a species in positive controls, “N” denotes its presence in negative controls, and “S” represents its presence in only a single replicate of all samples.

#### 2.3.4 Re-annotation with updated reference

MitoNGS has adapted a user-friendly strategy to facilitate re-annotation upon database updates. Unlike the legacy MiFish pipeline, which exclusively accepts raw NGS reads, MitoNGS accepts users-provided ASVs or Operational Taxonomic Units (OTUs) as input data for taxonomy annotation. During the initial analysis, MitoNGS generates annotation results from raw NGS reads and provides users with a unique identifier (job ID) and a downloadable file of ASV sequences. Within three months, users can utilize this job ID to retrieve ASVs and parameter settings for re-annotation. After three months, old job files would be removed from the server, but users can upload ASV sequence files to conduct re-annotation. Consequently, the time-consuming raw read processing and denoising step is bypassed during the re-annotation process, enhancing overall efficiency.

### 2.4 Testing datasets

Eight fish eDNA metabarcoding datasets were selected (Figure 5, Table 3), covering all the four widely used genes, 12S rRNA, 16S rRNA, cytochrome c oxidase I (COXI or COI), and cytochrome b (CYTB), and two main sequencing platforms (Illumina and Nanopore). The taxonomy identification results were compared with those in original publications. For the dataset of “Europa island in Western Indian Ocean, French Southern and Antarctic Lands” (Lamperti et al., 2024), OTU sequences were readily available and utilized as the initial input so that the result comparison would be easier to be conducted in a one-to-one relationship. For other datasets, raw reads were used to do the annotation.

**Figure 5.**
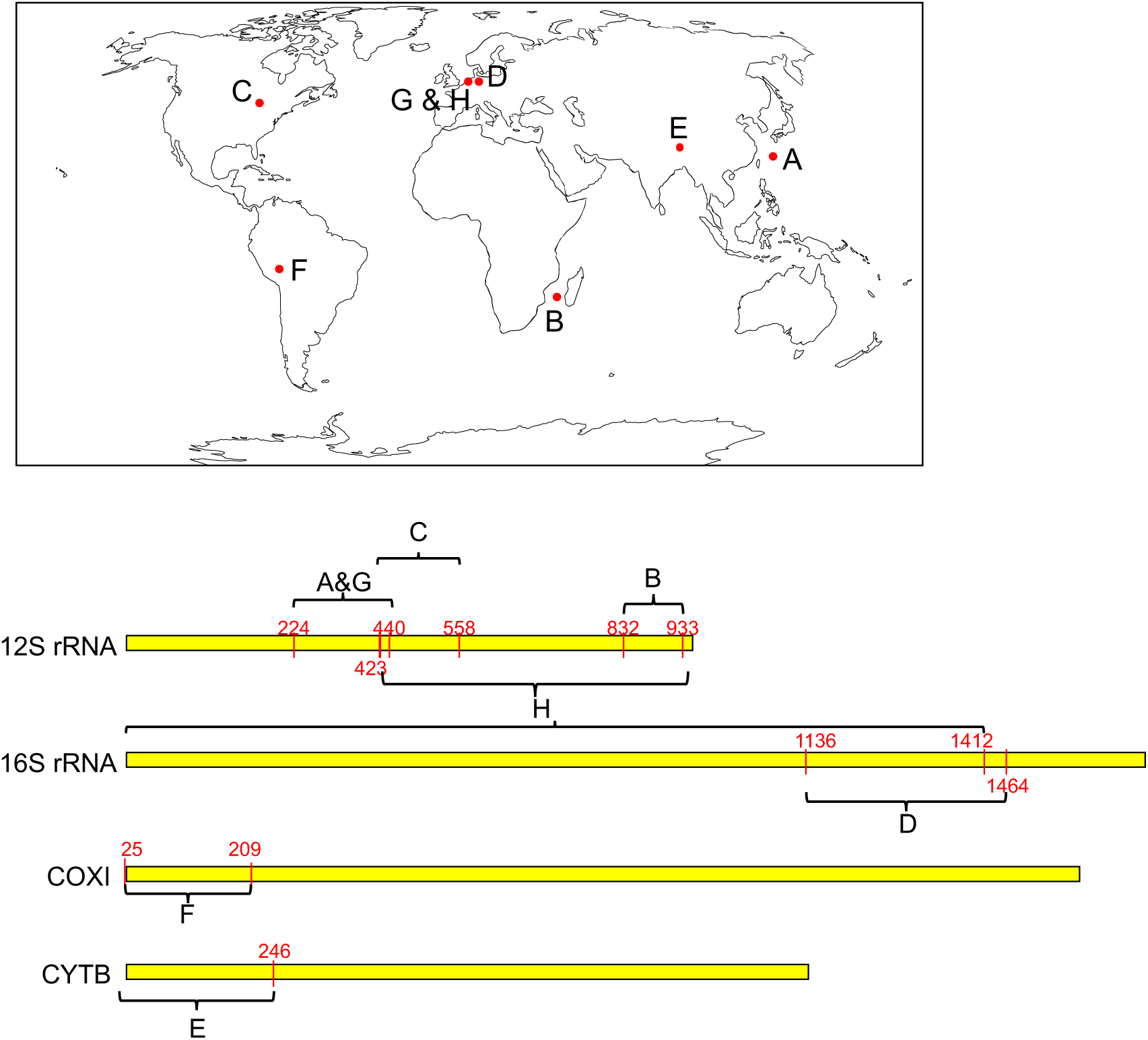
Sampling locations and marker regions of eight testing datasets. Coordinates of marker regions are based on the mitogenome of Japanese eel *Anguilla japonica* (AB038556). A) Uchidomari river in Okinawa Island, Japan (Sato et al., 2018). Marker is MiFish-U (Miya et al., 2015) in 12S rRNA. B) Europa Island in Western Indian Ocean, French Southern and Antarctic Lands (Lamperti et al., 2024). Marker is teleo (Valentini et al., 2016) in 12S rRNA. C) Boardman Lake in Michigan, USA (Gehri, Larson, Gruenthal, Sard, & Shi, 2021). Marker is Riaz (Kelly, Port, Yamahara, & Crowder, 2014; Riaz et al., 2011) in 12S rRNA. D) Mittelriede and Schunter River in Braunschweig, Germany (Vences et al., 2016). Marker is Vert-16S (Vences et al., 2016) in 16S rRNA. E) Yarlung Zangbo River in Tibet, China (Feng et al., 2023). Marker is customized by authors using the start region of CYTB. F) Madre de Dios region of the Peruvian Amazon, Peru (Timana-Mendoza et al., 2025). Marker is Mariac (Mariac et al., 2022) in COXI. G) North Sea Ray Reef aquarium in Harderwijk, the Netherlands (Doorenspleet et al., 2025). Marker is MiFish-U in 12S rRNA and sequenced by Nanopore. H) The same samples as G), while marker is customized by authors using a 2kb region covering 12S and 16S rRNA.

**Table 3.**
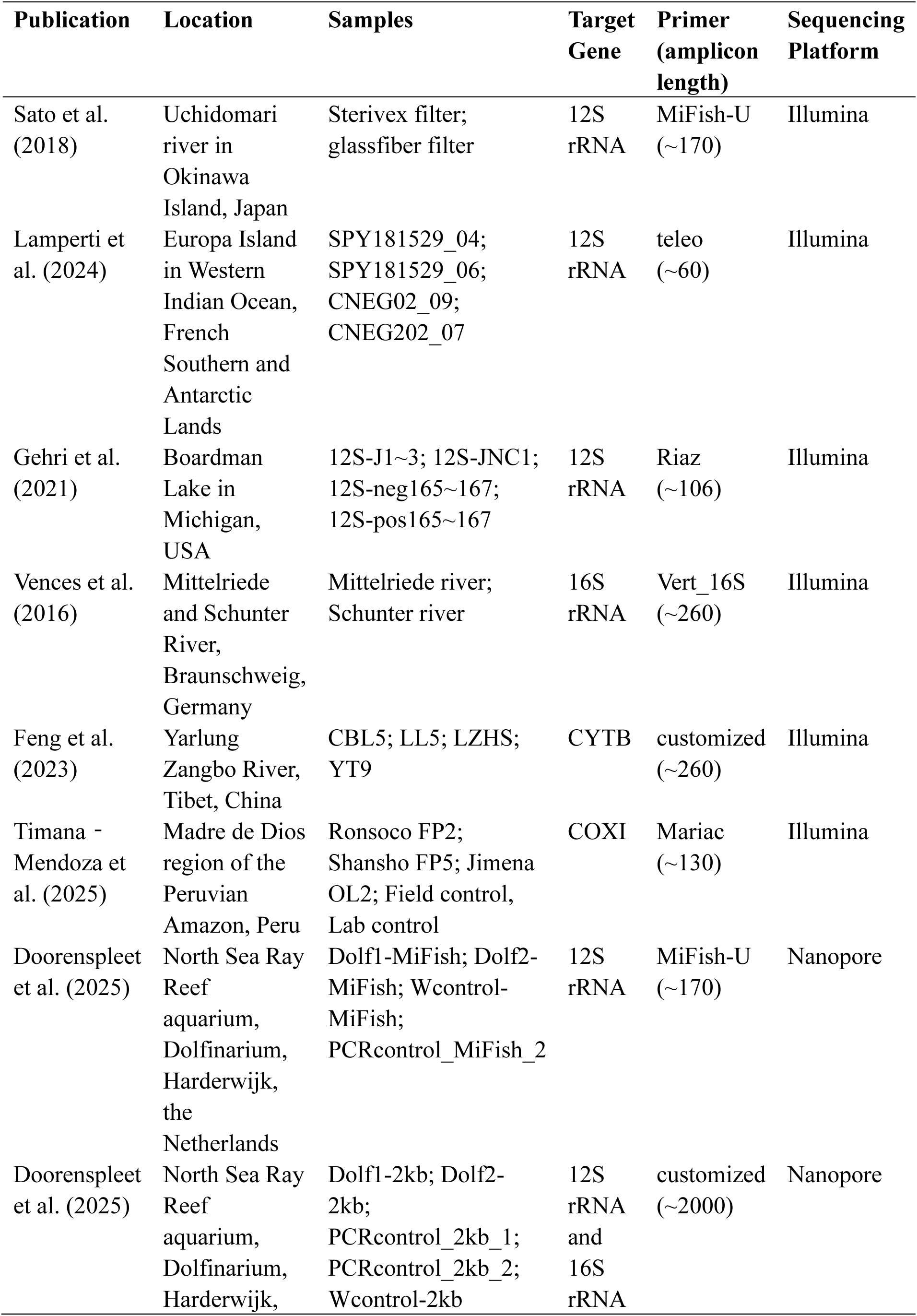
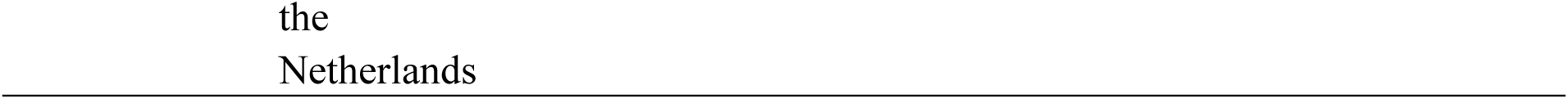
Eight datasets for testing the performance of MitoNGS.

| Publication | Location | Samples | Target Gene | Primer (amplicon length) | Sequencing Platform |
| --- | --- | --- | --- | --- | --- |
| Sato et al. (2018) | Uchidomari river in Okinawa Island, Japan | Sterivex filter; glassfiber filter | 12S rRNA | MiFish-U (~170) | Illumina |
| Lamperti et al. (2024) | Europa Island in Western Indian Ocean, French Southern and Antarctic Lands | SPY181529_04; SPY181529_06; CNEG02_09; CNEG202_07 | 12S rRNA | teleo (~60) | Illumina |
| Gehri et al. (2021) | Boardman Lake in Michigan, USA | 12S-J1~3; 12S-JNC1; 12S-neg165~167; 12S-pos165~167 | 12S rRNA | Riaz (~106) | Illumina |
| Vences et al. (2016) | Mittelriede and Schunter River, Braunschweig, Germany | Mittelriede river; Schunter river | 16S rRNA | Vert_16S (~260) | Illumina |
| Feng et al. (2023) | Yarlung Zangbo River, Tibet, China | CBL5; LL5; LZHS; YT9 | CYTB | customized (~260) | Illumina |
| Timana - Mendoza et al. (2025) | Madre de Dios region of the Peruvian Amazon, Peru | Ronsoco FP2; Shansho FP5; Jimena OL2; Field control, Lab control | COXI | Mariac (~130) | Illumina |
| Doorenspleet et al. (2025) | North Sea Ray Reef aquarium, Dolfinarium, Harderwijk, the Netherlands | Dolf1-MiFish; Dolf2-MiFish; Wcontrol-MiFish; PCRcontrol_MiFish_2 | 12S rRNA | MiFish-U (~170) | Nanopore |
| Doorenspleet et al. (2025) | North Sea Ray Reef aquarium, Dolfinarium, Harderwijk, the Netherlands | Dolf1-2kb; Dolf2-2kb; PCRcontrol_2kb_1; PCRcontrol_2kb_2; Wcontrol-2kb | 12S rRNA and 16S rRNA | customized (~2000) | Nanopore |

## 3 Results

### 3.1 Comprehensive reference database

#### 3.1.1 Coverage on species and clades

By June 2025, there were 905,179 sequences in the MitoFish database. Among them, 78.0% were protein-coding or rRNA sequences from 42,996 species, covering all 86 orders, 572 out of all 575 families, and 4,606 out of all 4,772 genera. Scytalinidae (graveldivers), Lacantuniidae and Hispidoberycidae are the only three families lacking mitochondrial sequences. These families are rarely studied, with fewer than 100 publications to date. Consequently, the reference database (MitoFish) is comprehensive, meeting the needs of global fish eDNA research.

Regarding the four widely used mitochondrial genes - 12S rRNA, 16S rRNA, COXI, and CYTB - we observed that while their sequences have covered all orders and most families, coverage on genera was lower (Table 4). Notably, for the most widely used 12S rRNA gene, sequences were available in only 77.8% of genera. This is understandable given that studies on 12S rRNA occurred later than those on COXI, which was initially used as a standard marker in many animals (Collins et al., 2019). This gap will be filled by the rapid increase in newly sequenced species in the coming years.

**Table 4.** Sequence coverage of four widely used mitochondrial genes in order, family and genus level.

| **Gene** | **Orders** | **Families** | **Genera** |
| --- | --- | --- | --- |
| 12S rRNA | 86 (100%) | 548 (95.3%) | 3,713 (77.8%) |
| 16S rRNA | 86 (100%) | 558 (97.0%) | 3,909 (81.9%) |
| COXI | 86 (100%) | 561 (97.6%) | 4,247 (89.0%) |
| CYTB | 86 (100%) | 539 (93.7%) | 3,600 (75.4%) |

#### 3.1.2 Heterospecific regions

Among the 706,818 sequences containing protein-coding or rRNA genes, 127,971 contained at least one heterospecific segment. A typical example is one of the full mitogenome (AP006035) of Indo-Pacific/Atlantic sailfish *Istiophorus platypterus* (Istiophoriformes: Istiophoridae) (Claver et al., 2023). We identified that heterospecific regions occur in the end region of 12S rRNA gene and the start region of the 16S rRNA gene with ≥99% similarity to multiple *Engraulis* species (Clupeiformes: Engraulidae). Metabarcoding markers located at the beginning of 12S rRNA or the end of 16S rRNA would function normally, while those located at the end of 12S rRNA or the beginning of 16S rRNA would be affected (Figure 6).

**Figure 6.**
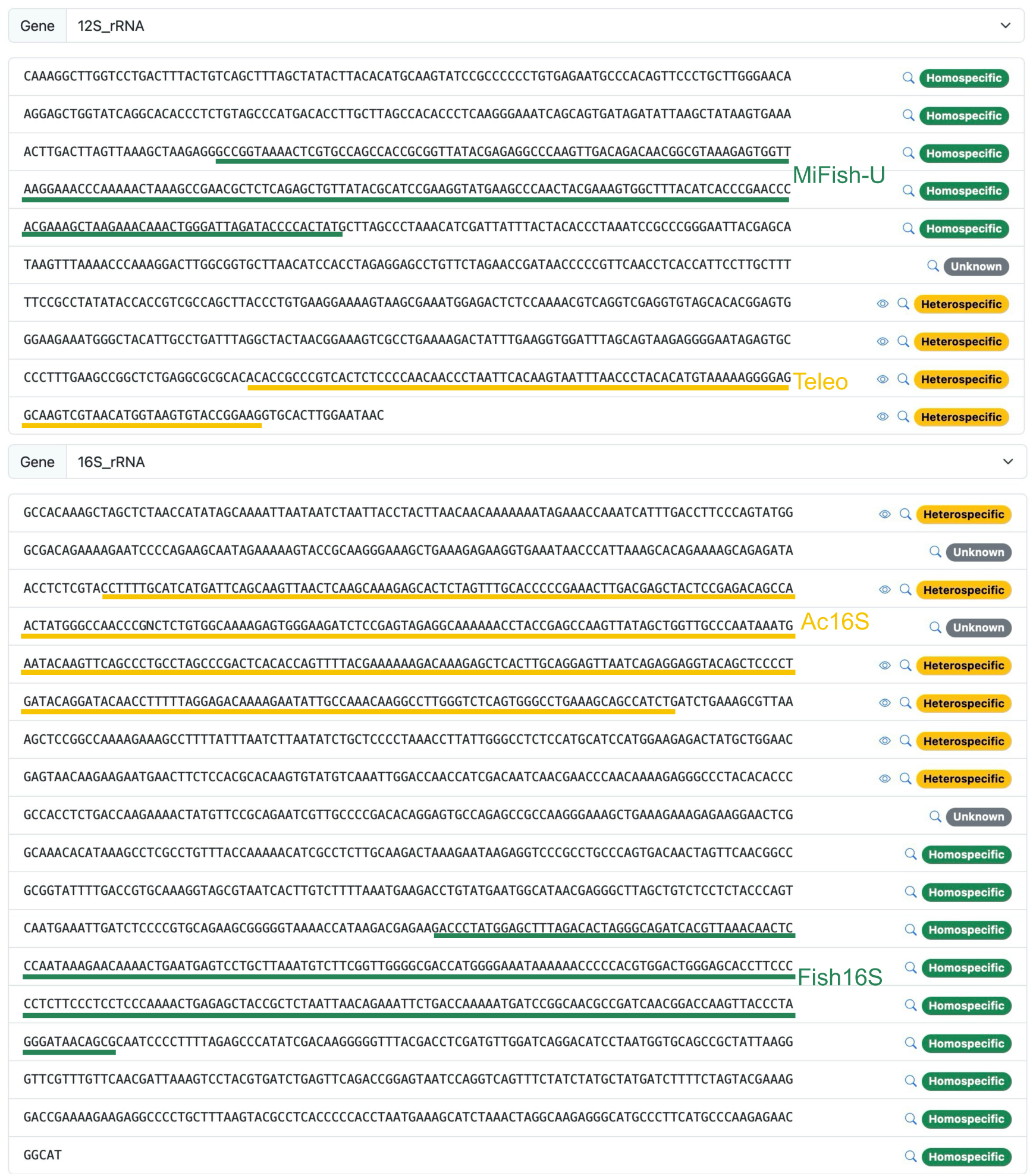
Heterospecific regions of AP006035, the mitogenome of Indo-Pacific/Atlantic sailfish *Istiophorus platypterus*. Amplicon regions of four commonly used metabarcoding primers were displayed: MiFish-U (Miya et al., 2015), Teleo (Valentini et al., 2016), Ac16S (Evans et al., 2016), and Fish16S (Berry et al., 2017).

#### 3.1.3 Environmental habitat and geographic occurrence data

Environmental habitat data and geographic occurrence data were retrieved from FishBase and GBIF, respectively, which primarily contained binomial species. Among the 42,996 species with protein-coding or rRNA sequences in the reference database, there are 22,810 binomial species, among which 22,075 (96.8%) are available with environmental habitat data from FishBase, and 21,247 (93.1%) with occurrence data in GBIF. As expected, non-binomial species seldom have environmental habitat data (0.17%) since they are not included in FishBase. Conversely, the proportion of non-binomial species with occurrence data was higher (15.9%) because GBIF also includes occurrence records on DNA-based evidence.

### 3.2 Improved metabarcoding results in species-level resolution

#### 3.2.1 The benefit of comprehensive reference database

MitoNGS yields consistent results across all the eight testing datasets (Table 5). Notably, 32 cases were identified where ASVs or OTUs initially annotated at higher taxonomic levels were subsequently re-annotated to the species level (Supplementary Table S1C, S2D, S3C, S4C, S7C, S8C). One notable case is a 66bp-OTU sequence from the dataset of Europa Island in Western Indian Ocean (Lamperti et al., 2024). Initially annotated as a superclass “Actinopterygii” (Supplementary Table S2D), the sequence underwent re-annotation to the species *Caracanthus unipinna* (Perciformes: Caracanthidae) through MitoNGS. The reference sequence (LC685936) of the best match was published in February 2022, later than the reference database (March 2020) used in the original publication. It was the only one with high similarity of the OTU sequence, and the species *Caracanthus unipinna* has occurrence records near the sampling location (Figure 7), corroborating the accuracy of the re-annotation. For cases where species (or higher taxa) were different from the original results, the majority of them were also improved due to higher matching similarity in the updated references (Supplementary Table S1D, S2E, S3E, S7E, and S8D). This demonstrates the significant utility of employing the most recent and comprehensive reference database in achieving high-resolution taxonomy identification outcomes.

**Figure 7.**
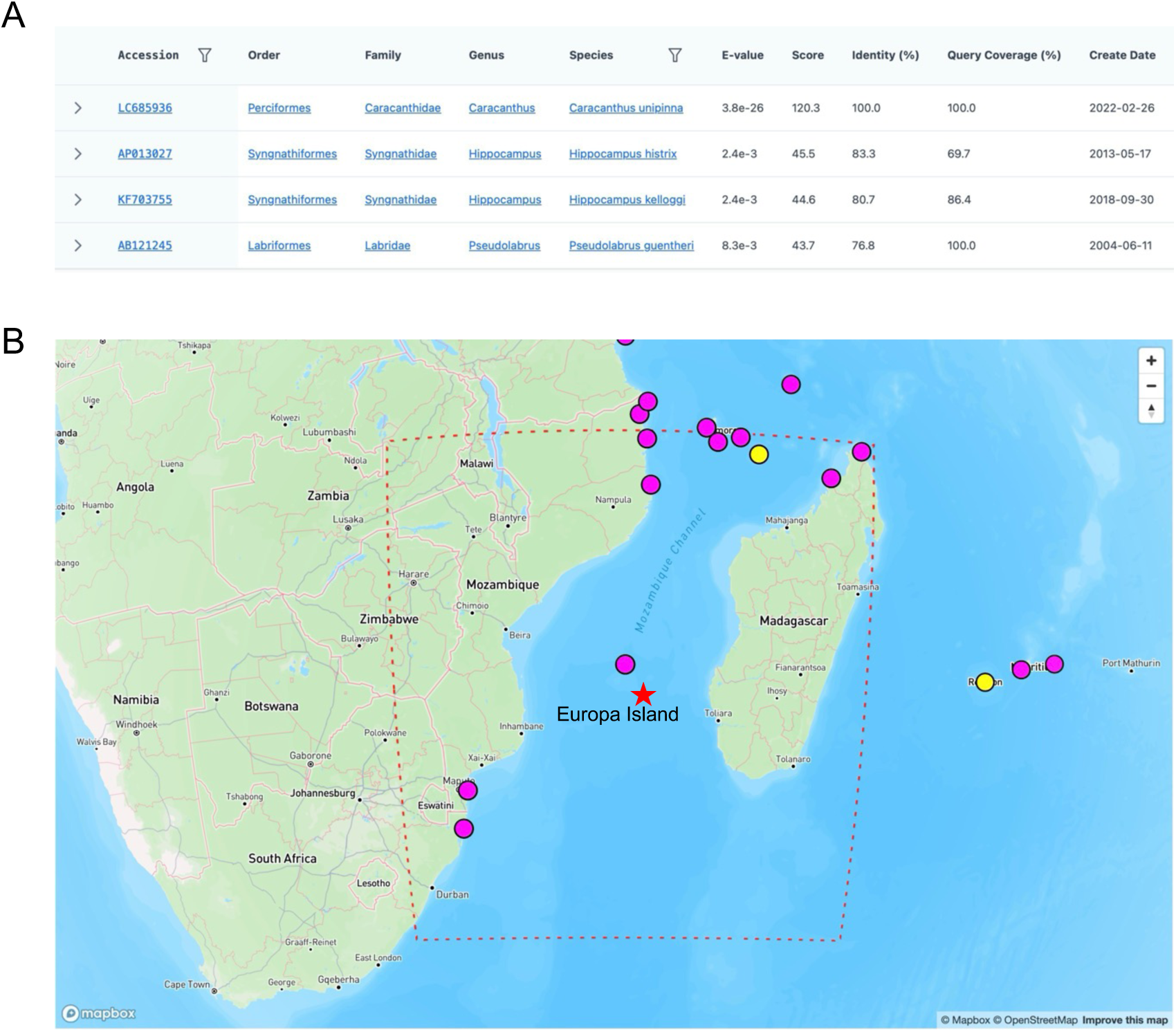
Example of higher taxon (superclass: Actinopterygii) reannotated in species-level (*Caracanthus unipinna*). The 66bp-OTU (CCCCGATACATCACCACTCGTATATTAACAAGCCTAGATAAAATACTGAGGGGAGGCAAGTCGTAA) was from the dataset of Europa Island in Western Indian Ocean (Lamperti et al., 2024). A). BLAST results (screenshot via MitoFish at https://mitofish.aori.u-tokyo.ac.jp/blast/simple) of the OTU to top four hits. Similarity of all other three hits are lower than 85%. B) The position of the sampling site Europa Island (marked with star) and the occurrence records of *Caracanthus unipinna* (screenshot via MitoFish at https://mitofish.aori.u-tokyo.ac.jp/detail/species/1146959). Pink points indicate observation of specimen materials (such as those used for sequencing) while yellow points indicate observation of live creatures.

**Table 5.**
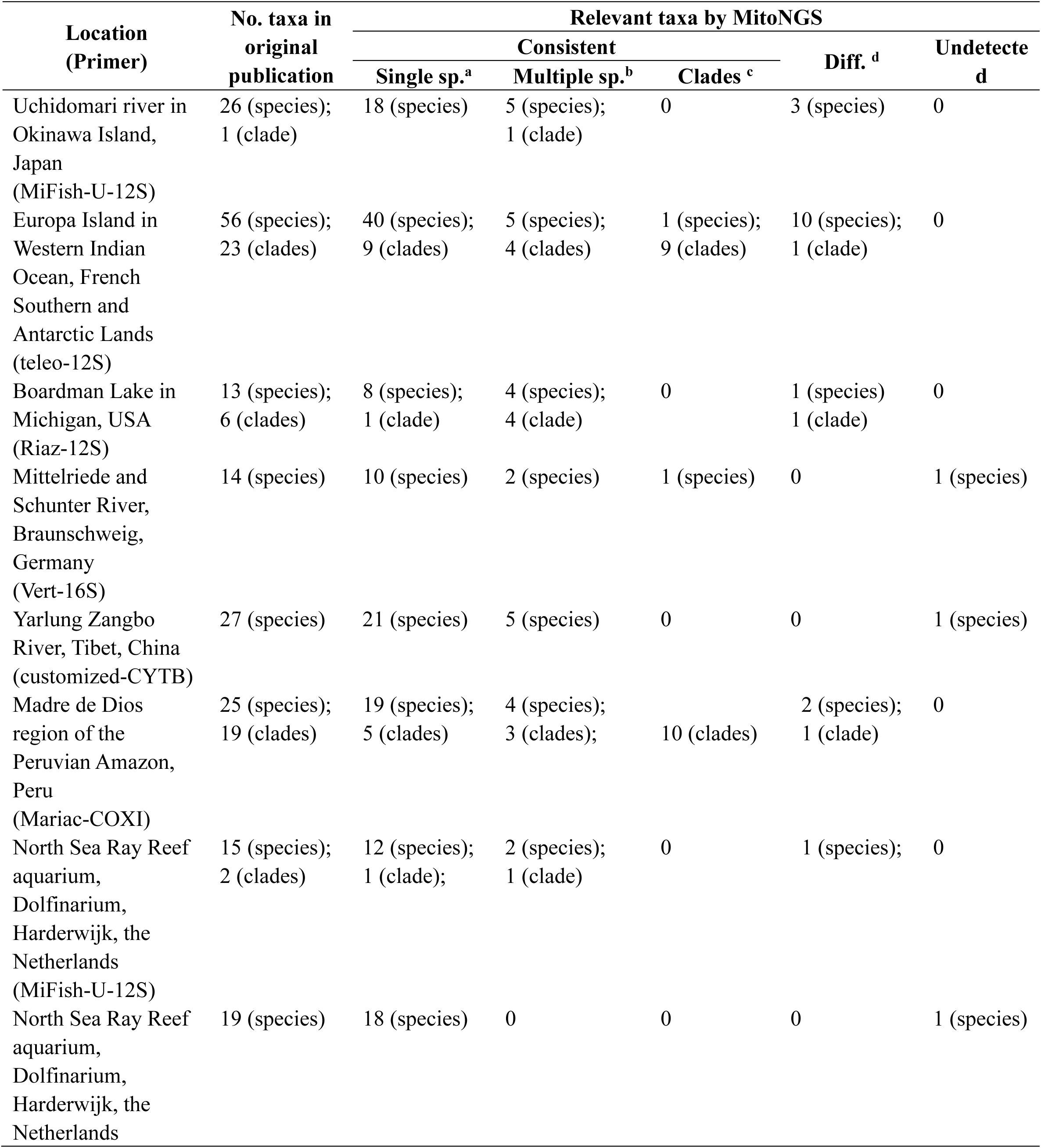
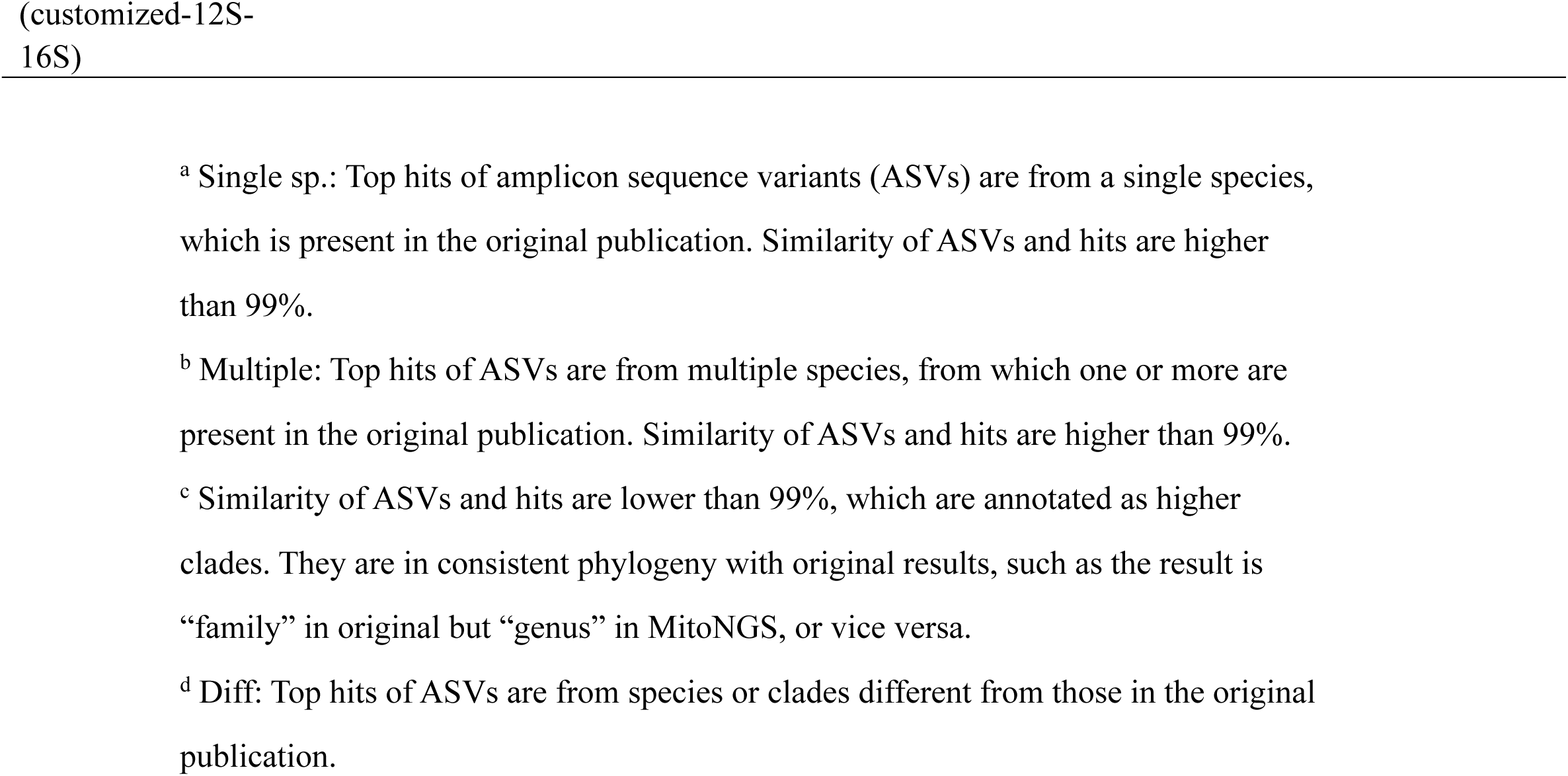
Comparison of taxonomy identification results of MitoNGS and the original literature. See Supplementary Table S1∼9 for the full lists of taxa.

#### 3.2.2 Species groups and further filtration using heterospecific information

Except for the only dataset with long amplicons (2Kb), all the remaining seven datasets include species groups where multiple species share identical metabarcoding regions (Table 5). It is reasonable as short markers inherently exhibit reduced distinguishability. In the result table, environmental habitat data and geographic occurrence data were presented to facilitate further filtering, and all related reference sequences of these species could be downloaded (Figure 8).

**Figure 8.**
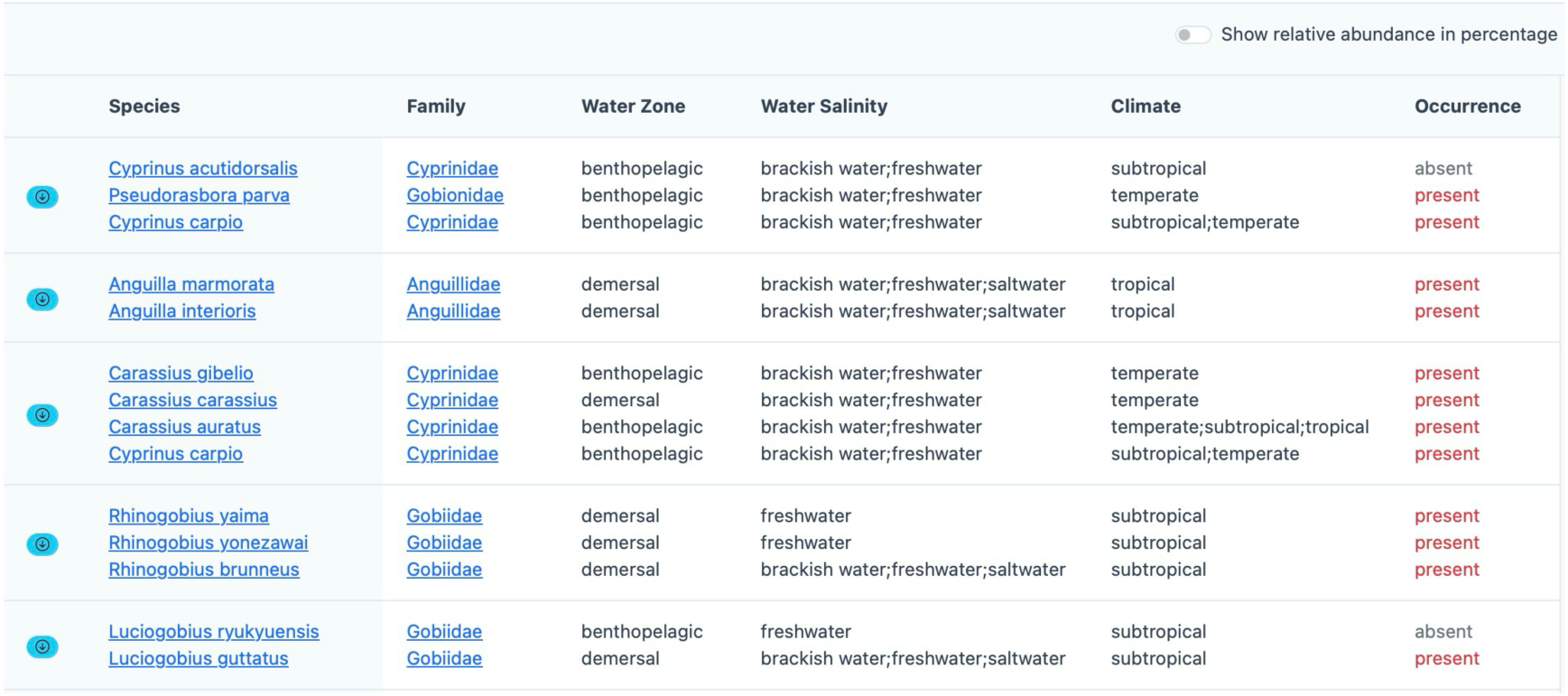
Species group detected in MitoNGS. Screenshot from the dataset of Uchidomari river in Okinawa Island, Japan (Sato et al., 2018). Species sharing identical ASVs are displayed in the same row, with environmental habitat and occurrence data shown after. The attached download button in the leftmost column enables users to download all the mitochondrial sequences of these species to conduct further analysis, such as designing species-specific qPCR primers.

As indicated in Figure 3, heterospecific information is employed to facilitate the filtering of species groups. We compared the results with those that did not apply this filtration and indeed identified 10 cases where unambiguous results were obtained by heterospecific filtration (Supplementary Table S9). In these cases, ASVs have top hits from multiple species, among which only one species is verified to be non-heterospecific, demonstrating that the inclusion of heterospecific information does indeed contribute to the enhancement of taxonomy resolution.

### 3.3 Discovery of novel fish species and off-target amplifications

15 novel fish species were detected in three datasets (Table 6, Supplementary Table S1E, S6C, S7E, and S9B). Among these, 10 species have occurrence records near the sampling regions, indicating true positive discoveries. The remaining five species lacking occurrence evidence might be invasive ones.

**Table 6.**
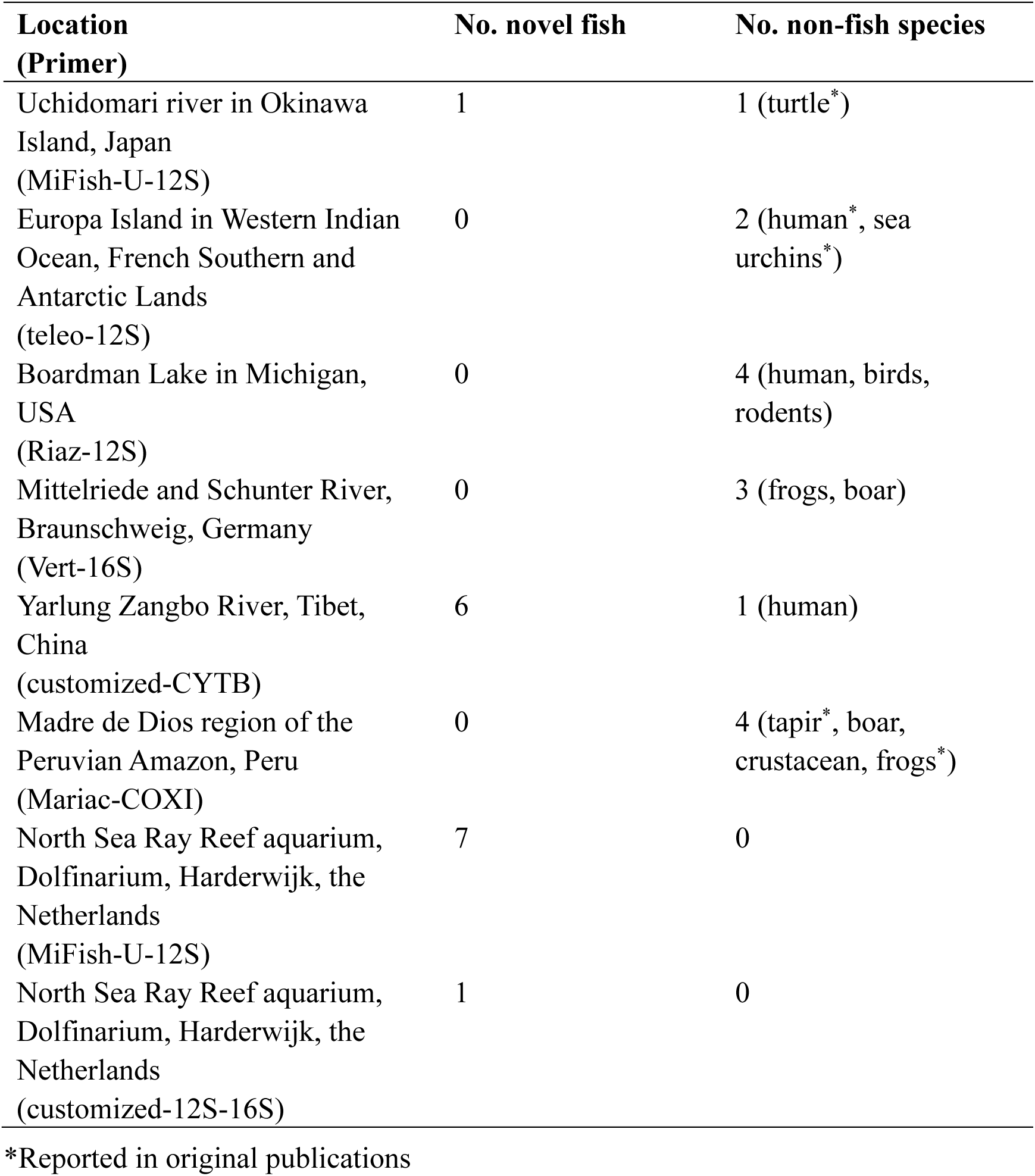
Novel fish species and off-target products detected by MitoNGS. Only species satisfying all the following criteria were counted. 1) unambiguous; 2) identities higher than 99%; 3) not detected in negative controls; 4) detected in at least two replicates if available; 5) relevant BLAST hits not in the heterospecific region (for fish species).

Non-fish species with identity ≥ 99% were detected in six datasets (Table 6, Supplementary Table S1A, S2A and E, S3F, S4D, S5D, and S6F), encompassing a range of taxa such as invertebrates, amphibians, and mammals. In fact, non-fish taxonomies were also detected in the remaining two datasets, albeit with a lower identity or only present in a single replicate within samples, potentially indicating contamination. Unassigned ASVs (Figure 3) are absent from all the eight datasets.

## 4 Discussion

### 4.1 Excellent performance

In all testing datasets, taxonomy identifications are consistent with original ones, while most inconsistent cases were actually improved annotations due to our more comprehensive reference database. Notably, the analysis speed is fast, particularly for Nanopore reads. Typically, generating consensus sequences from raw Nanopore reads is time-consuming. In our testing datasets, the Dolf2-2kb library from North Sea Ray Reef aquarium in the Netherlands (Table 3) contains 141,302 long reads with an average length of 2kb. MitoNGS completed the entire analysis using 48 threads on our server in approximately 6 minutes. In contrast, the DECONA pipeline used in the original publication (Doorenspleet et al., 2025) required over 1 hour to run using 60 threads. In summary, MitoNGS achieves a favorable cost-performance ratio by balancing the speed and accuracy.

Although there were five undetected cases (Table 6), all of them were plausible due to insurmountable obstacles. *Cyprinus carpio* in the dataset of Mittelriede River, Germany (Supplementary Table S4A) was in low abundance (9). Although the abundance of *Triplophysa stolickai* in the dataset of Yarlung Zangbo River, Tibet, China (Supplementary Table S5A) was higher (66 in sample LZHS), we failed to detect any sequences in high similarity (≥ 98% based on the authors’ methods) with *Triplophysa stolickai*. Regarding *Chelidonichthys lucerne* in the Nanopore 2Kb libraries (Supplementary Table S8A), we found that *Chelidonichthys lucerne* lacks long reference sequences by June 2025. The longest rRNA gene, ON000326, was 1,126bp, resulting in only ∼50% query coverage of the ∼2kb Nanopore reads, and was therefore not included in the final results. Except for the dataset of Europa Island in French Southern Lands (Lamperti et al., 2024), ASVs’ (or OTUs’) sequences were not publicly available for the remaining seven datasets, making the one-to-one comparison difficult. It would be beneficial for all metabarcoding-related research to make amplicon sequences publicly available, as they are computationally intensive but reference-independent. This would facilitate downstream validations, particularly on updated references.

### 4.2 Reference gaps

Fish ASVs with similarities below 99% were detected in all the eight datasets, suggesting the presence of potentially unknown fish species not included in the reference database. To further investigate, in each dataset, we manually inspected five ASVs with the lowest similarities by searching them again in GenBank. The results indicated that for COXI metabarcoding, ASVs with extremely low similarities (< 80%) might originate from bacteria COXI genes (similarities ∼ 90%). Given the limited data resources for accurate bacteria COXI genes, they were not included in the background database. This omission presents an opportunity for future efforts in MitoFish and MitoNGS. For all other genes, no novel hits were found in GenBank, suggesting that a plenty of fish species are yet to be discovered and/or sequenced.

### 4.3 Strategies to improve taxonomy resolution

For ASVs with identical top-hits from multiple species, MitoNGS displays species groups instead of LCA. Among all the species groups detected in our testing datasets, except for genus *Gomphosus* (Supplementary Table S2B) and *Pomoxis* (Supplementary Table S5B), which only contains two species respectively, all other clades include additional species that do not belong to the species groups. This demonstrates that species groups provide a more precise representation of ASVs’ taxonomy compared to LCA.

Furthermore, MitoNGS employs binomial status and heterospecific information to filter species groups (Figure 3). Additionally, environmental habitat and geographic occurrence data within groups are also displayed (Figure 8). MitoNGS does not automatically exclude species lacking occurrence records near sampling locations, as occurrence data is often incomplete. For instance, in the testing dataset of Uchidomari river, Okinawa, Japan (Supplementary Table S1C), *Luciogobius guttatus/ryukyuensis* were detected. Although the latter lacked occurrence records with geographic coordinates, there were 91 records in GBIF with the area roughly labelled as “Japan” by the end of June 2025, suggesting that *Luciogobius ryukyuensis* could still be the correct identification. There are no universally accepted guidelines for whether to retain or exclude species with varying habits or occurrence patterns, and this decision should depend on the research objectives and prior knowledge of sampling environments.

We also observed that certain species within some groups could form interspecific hybrids (Supplementary Table S1B, S1D, S2D, S3B, S3C and S5B), indicating insufficient reproductive barriers. From another perspective, such species groups could be considered virtual species. Currently, data on interspecific hybrids and their evidence levels are scarce. In the future, with the accumulation of hybrid cases, species based on genuine reproductive barriers could be constructed to serve as the basic unit of metabarcoding results.

Long-read-based metabarcoding demonstrated exceptional performance in resolving closely related species. In our testing dataset, a 2kb Nanopore library successfully identified all the species unambiguously (Supplementary Table S8). The primary challenge lies in the scarcity of long reference sequences, which will be addressed with the accelerated accumulation of complete mitogenomes.

## 5 Conclusion

MitoNGS is an integrated, web-based platform for the analysis of fish metabarcoding data across diverse mitochondrial loci. By combining curated reference databases, heterospecific region annotation, off-target filtering, environmental metadata, and species occurrence information, MitoNGS enables high-resolution, species-level taxonomic assignments. As the successor to the MiFish pipeline, it addresses several longstanding challenges in eDNA analysis, including ambiguous barcodes and limited marker compatibility. With its flexible input formats and support for long-read sequencing technologies, MitoNGS is poised to advance the field of aquatic biodiversity monitoring and support a wide range of ecological, conservation, and applied research efforts.

## Notes

https://mitofish.aori.u-tokyo.ac.jp/mito-ngs

### Competing Interest Statement

The authors have declared no competing interest.

### Funding

- The University of Tokyo: OceanDNA Project (No. 027) from UTokyo Future Society Initiative, Interdisciplinary Collaborative Research Program (1-B) of Atmosphere and Ocean Research Institute
- Ministry of Education, Culture, Sports, Science and Technology, Japan

